# Type 1-polarized DC immunotherapeutic contains heterogeneous populations with IL-12p70 production restricted to a rare subset

**DOI:** 10.64898/2026.04.29.721620

**Authors:** Allison E. DePuyt, Peter E. J. Shoucair, Holly Bilben, Jeremy J. Martinson, Bernard J. C. Macatangay, Charles R. Rinaldo, Pawel Kalinski, Robbie B. Mailliard

## Abstract

Monocyte-derived DC therapies programmed for robust IL-12p70 production have been associated with favorable outcomes in cancer clinical trials. However, clinical responses remain inconsistent even under standardized protocols, and the cellular basis for this variability is unknown. We leveraged single-cell multiomics to characterize two widely used DC platforms, the high-IL-12p70-producing alpha-Type-1-polarized DC (αDC1) and the IL-12p70-deficient DC induced in the presence of PGE_2_ (PGE_2_-DC), at baseline and following rhCD40L activation. DC generated from 7 healthy participants representing the spectrum of rhCD40L-induced IL-12p70 production were profiled by transcriptome analysis with concurrent 42-plex surface proteomics, multiplex ELISA, and ELISpot quantification of IL-12p70-producing cells. While αDC1 and PGE_2_-DC distinctly responded to rhCD40L, αDC1 alone unexpectedly comprised 3 transcriptionally and phenotypically distinct subpopulations in resting and stimulated states. Only a limited fraction of αDC1 coexpressed *IL12A* and *IL12B* (IL-12p70 producers), which was confirmed by ELISpot at the protein level. The distribution of αDC1 subclusters varied markedly between individuals and correlated with bulk cytokine and chemokine secretion profiles. Heterogeneity within αDC1 preparations may underlie inconsistent clinical trial outcomes, and identification of associated surface proteins provides a prospective strategy for subcluster enrichment to enhance DC release criteria and patient stratification for optimized therapeutic efficacy.

## INTRODUCTION

DC-based immunotherapies are a promising treatment strategy for both cancers and infectious diseases (1,2), leveraging their unique capacity to prime and coordinate appropriate antigen-specific T cell responses. The clinical potential of this approach was underscored by the FDA approval of sipuleucel-T for metastatic hormone refractory prostate cancer, which prolonged overall survival and was well-tolerated in clinical trials (3,4). Despite this landmark achievement, DC therapy efficacy has remained inconsistent across clinical applications. Trials have utilized diverse DC preparations with distinct functional properties, including monocyte-derived DC, peripherally circulating CD1c^+^ myeloid DC, and activated peripheral blood mononuclear cells (5). While vaccination with natural CD1c^+^ DC induced durable CD8^+^ T cell responses and improved survival in some metastatic melanoma patients (6), a recent trial using similar DC failed to demonstrate a benefit in recurrence-free survival despite the development of antigen-specific responses (7). Because autologous monocyte-derived DC are routinely generated and matured ex vivo to differentially express certain functional traits, a wide range of strategies have been developed for their specific production and clinical application.

One widely adopted protocol for generating clinically applicable DC combines a cocktail of inflammatory cytokines along with prostaglandin E_2_ (PGE_2_) to yield a mature DC (PGE_2_-DC) with favorable lymph node homing and robust surface expression of costimulatory molecules essential for inducing primary T cell responses (8). These PGE_2_-DC, however, have a reduced capacity to produce IL-12p70 (9), a key factor in DC-mediated Th1 polarization of CD4^+^ T cells and optimal priming of cytotoxic CD8^+^ lymphocytes (CTL) (10,11). PGE_2_-DC are particularly weak in their responsiveness to CD40L, a helper signal provided by antigen activated CD4^+^ T cells that is critical to supporting CTL responses (12–16). The alpha-type-1 polarized DC (αDC1) platform, which substitutes PGE_2_ with type-I and -II interferons combined with polyinosinic:polycytidylic acid (poly(I:C)), was therefore specifically designed for optimal IL-12p70 production in response to the T helper signal CD40L (15). In addition to their superior IL-12p70 production, αDC1 demonstrate enhanced MHC-I antigen presentation and cross-presentation (17), produce chemokines that promote more effective interactions with naïve and effector T cells, and display a unique capacity to mediate intercellular antigen transfer via CD40L-induced tunnelling nanotube formations (18).

Despite these mechanistic insights and protocol optimizations, clinical outcomes remain variable even among patients receiving αDC1 vaccines within the same trial protocol. Importantly, multiple trials have indeed supported the clinical relevance of DC vaccine-derived IL-12p70: IL-12p70 production by an αDC1 therapeutic positively correlated with time to disease progression in patients with recurrent malignant glioma (19), and neoantigen-loaded IL-12p70-secreting DC induced robust tumor-specific CTL responses in patients with melanoma (20). Moreover, in a recent Phase II trial treating checkpoint-refractory advanced melanoma, αDC1 therapeutics from patients with positive clinical outcomes (6 out of 13 treated) displayed increased bulk expression of genes associated with antigen processing and interferon/toll-like receptor signaling compared to those from non-responders (21). The presence of key costimulatory proteins analyzed as part of typical quality control did not differ between these two patient groups (21), suggesting that functionally relevant heterogeneity exists within these DC preparations that appear uniform by standard phenotyping. This finding, in combination with the incomplete understanding of factors governing IL-12p70 production by individual DC (22), underscores the importance of investigating this heterogeneity to improve therapeutic efficacy.

Single-cell technologies have begun to uncover unexpected variety within endogenous DC populations previously considered homogeneous. Single-cell RNA sequencing of human peripheral blood revealed novel DC subtypes with distinct functions (23). It is therefore plausible that therapeutic DC preparations may also consist of functionally distinct subpopulations; however, the clinically relevant αDC1 and PGE_2_-DC platforms have yet to be interrogated with single-cell multiomics. In this study, we leveraged single-cell RNA sequencing combined with surface protein analysis to characterize heterogeneity within αDC1 and compare this as well as their CD40L response to that of conventional PGE_2_-DC. We demonstrate that αDC1, but not PGE_2_-DC, consist of transcriptionally and phenotypically distinct subsets with unique cytokine and chemokine profiles, with only a limited fraction of the cells producing a relatively large amount of IL-12p70 in response to CD40L. These subpopulations include cells biased toward Th1-promoting, Th2/T_reg_-attracting, and Th17-promoting responses. Importantly, the relative distribution amongst these subclusters varies between individuals. We also identify surface markers distinguishing these αDC1 subpopulations at baseline (prior to CD40L exposure), providing a strategy for their isolation or patient stratification to optimize therapeutic efficacy.

## RESULTS

### αDC1 production of IL-12p70 is limited to a minor population of CD40L-responsive cells

Though αDC1 and PGE_2_-DC have been widely used in clinical trials to treat cancers and infectious diseases (1,2), modern single-cell multiomics methods have yet to be applied to deeply characterize these platforms and gain mechanistic insights into their functional disparities. To investigate the key differences between αDC1 and PGE_2_-DC responses to CD40L stimulation, we first differentially matured human primary monocytes according to two previously established protocols (8,15). Briefly, the pro-inflammatory αDC1 were generated using a combination of type-I and type-II interferons along with poly(I:C), a double-stranded RNA analogue. The PGE_2_-DC were matured using a standard cocktail of factors including prostaglandin E_2_, a lipid mediator of chronic inflammation. Both αDC1 and PGE_2_-DC highly express the maturation markers CD83 and CD86; however, αDC1 uniquely express Siglec-1/CD169 and PGE_2_-DC express OX40L (Figure 1A, B). Furthermore, αDC1 characteristically displayed amplified production of IL-12p70 in response to rhCD40L stimulation while PGE_2_-DC exhibited a blunted response (Figure 1C). Though this well-established increase in induced IL-12p70 production by αDC1 (15) was consistently observed within individual participants, the absolute amount of IL-12p70 released varied widely between individuals (Figure 1C). To determine whether this individual variation was due to differences in the number of IL-12p70-producing cells present or in the amount of IL-12p70 being released on a per cell basis, we next assessed the CD40L-activated DC preparations by IL-12p70 ELISpot assay. Unexpectedly, and despite clear spot development, only 0.57% of αDC1 produced IL-12p70 when treated with rhCD40L for 24 hours (Figure 1D, E). We also stimulated the αDC1 using CD40L-transfected J558 cells (CD40L-J558) to provide a stronger signal from membrane-associated CD40L. While CD40L-J558 stimulation increased the percentage of detectable IL-12p70-secreting αDC1 to 1.84% (Figure 1D), this was still much lower than originally anticipated.

**Figure 1.**
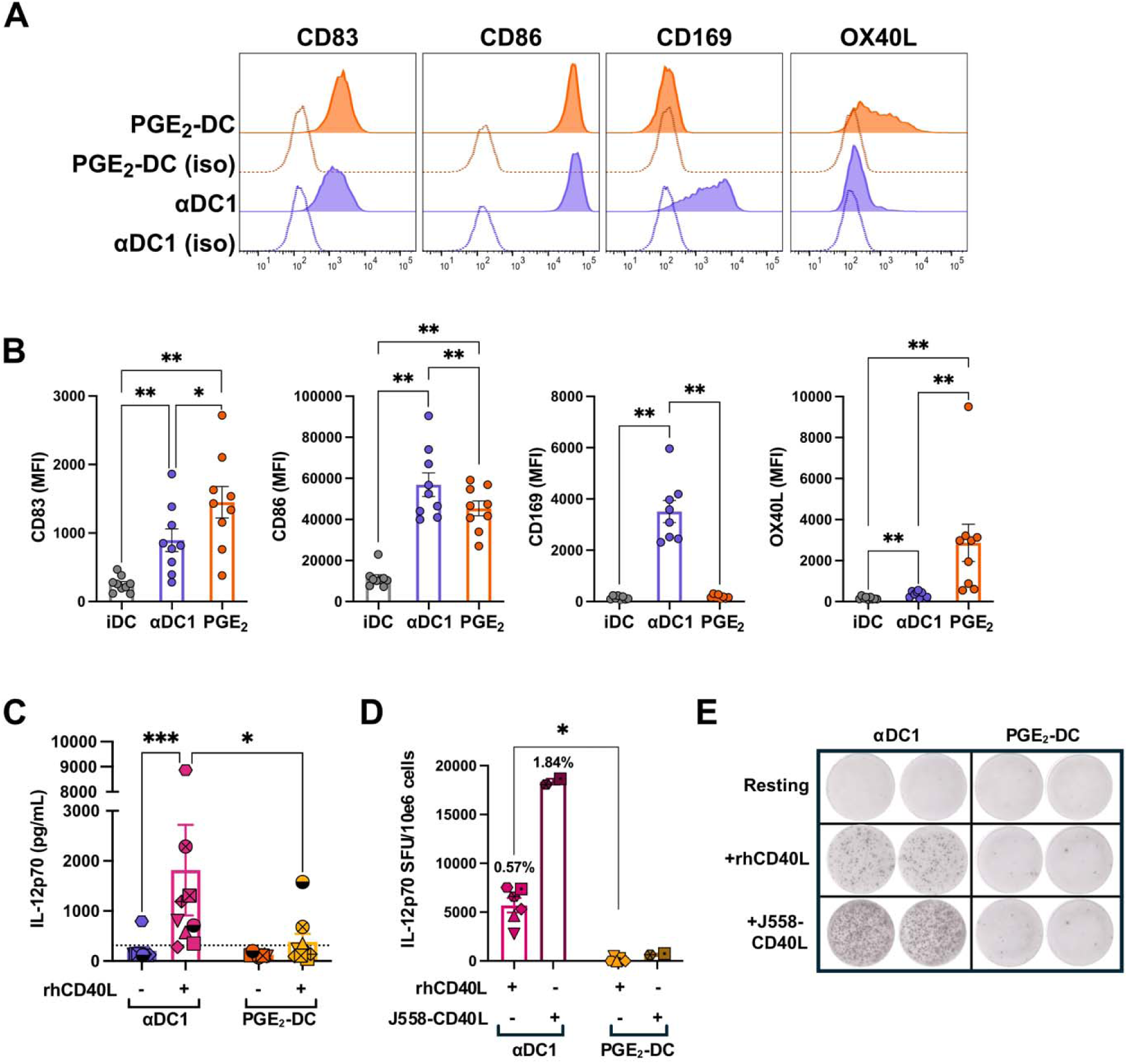
IL-12p70 production by αDC1 is limited to a minor subpopulation of CD40L-responsive cells. **A)** Differentially matured DC were analyzed for surface expression of CD83, CD86, CD169, and OX-40L by flow cytometry (representative *n* =1; summary of all experiments in panel B). Peaks with dashed lines represent isotype controls and filled peaks with solid lines indicate positive staining. **B)** MFI values for these surface markers on immature DC (iDC), αDC1, and PGE_2_-DC. *P* values were determined within each marker by paired Wilcoxon signed-rank tests with Bonferroni correction (*n* = 10). Bars represent mean ± SEM. **C)** Mature DC were stimulated with rhCD40L (1 µg/mL) for 24 hours to measure their bulk IL-12p70 production capacity. Data are normalized to cellular ATP content and presented as pg/mL of IL-12p70 per 200,000 cells. Bars represent mean ± SEM, and dashed line indicates the lower limit of detection. *P* values were determined by repeated measures two-way ANOVA with Šídák’s multiple comparisons test on log-transformed data (*n* = 9). **D)** The frequency of IL-12p70-secreting cells among rhCD40L- or J558-CD40L-stimulated αDC1 and PGE_2_-DC was determined by ELISpot assay and expressed as spot-forming units (SFU) per 10⁶ cells. Labels indicate the calculated mean proportion of IL-12p70-producing cells (*n* = 9) and bars represent mean ± SEM. *P* values were determined by paired Wilcoxon signed-rank test. **E)** Representative well images from the IL-12p70 ELISpot show spot formation indicating IL-12p70 release. In all bar plots, individual participant values are shown with unique symbols. For all statistical comparisons, \**p* < 0.05, \*\**p* < 0.01, \*\*\**p*<0.001.

### αDC1 and PGE_2_-DC differentially respond to CD40L

Given the surprising finding that only a percentage of αDC1 produced IL-12p70 when stimulated with rhCD40L, we utilized whole transcriptome scRNA-seq to determine the response status of the non-IL-12p70-producers and to evaluate the CD40L responsiveness of the two DC platforms on a broader scale. Importantly, the αDC1 and PGE_2_-DC each displayed unique transcriptional profiles at baseline as well as upon rhCD40L stimulation, with the clusters clearly separated in the UMAP by both DC type and stimulation status (Figure 2A; Supplemental Figure S1). Notably, for each DC type, the entire population of cells were responsive to rhCD40L signaling. In total, αDC1 differentially expressed 1531 unique genes that PGE_2_-DC do not in response to CD40L (Figure 2B). This gene set included upregulation of multiple chemokines such as *CCL3*, *CXCL10*, and *CCL4* that are known to attract various inflammatory cells including Th1, Th17, natural killer, and CD8^+^ T effector cells (Figure 2C) (24,25). Conversely, PGE_2_-DC increased expression of multiple genes associated with a tolerogenic phenotype (*LRAT, RBP4, SIGLEC15,* and *MAFB*) when stimulated with rhCD40L (Figure 2C) (26–28). Direct comparison of the CD40L-treated DC types identified a similar pattern: pathway enrichment analysis of αDC1 identified enriched terms indicating strong pro-inflammatory T cell priming ability, including antigen processing and presentation, human cytomegalovirus infection, and Fc-gamma R-mediated phagocytosis (Figure 2D, E), while PGE_2_-DC were enriched for pathways suggesting a suppressive transcriptional program, including Wnt signaling and the Polycomb repressive complex (Fig 2D, F) (29–31).

**Figure 2.**
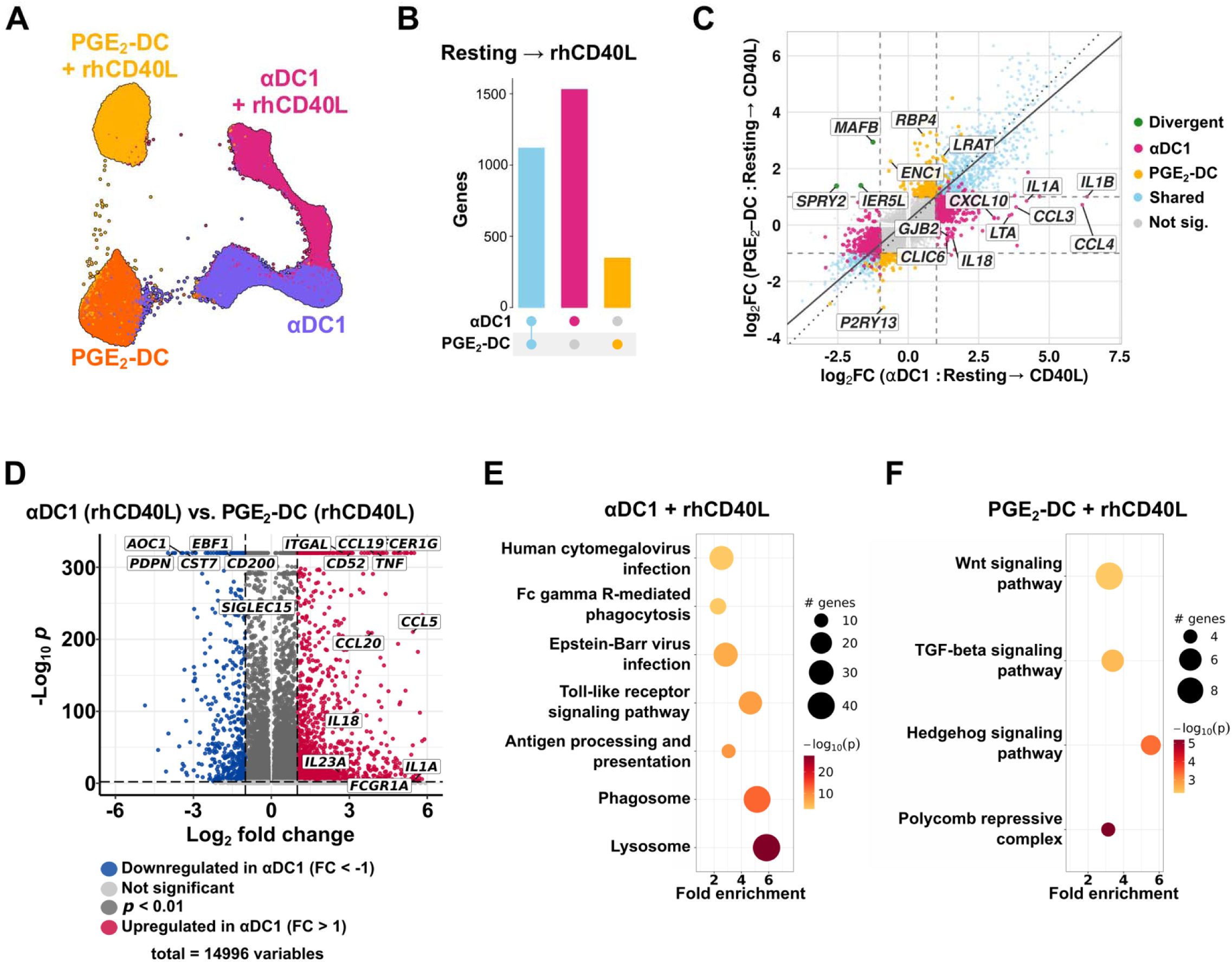
αDC1 mount a transcriptionally distinct CD40L response enriched for antigen processing and type-1 immune pathways, and display heterogeneity at baseline and following CD40L stimulation. **A)** UMAP of whole transcriptome single-cell RNA sequencing data from resting and rhCD40L-treated (4 hours) αDC1 and PGE_2_-DC (*n* = 7 participants; *n* = 149,535 cells). Clusters are colored by DC type and stimulation. **B)** UpSet plot depicting the number of differentially expressed genes (DEGs; adjusted *p* < 0.01 and |log_2_FC| > 1) induced by 4 hours of rhCD40L treatment in αDC1 (pink) and PGE_2_-DC (orange). Intersecting DEGs are represented by the blue bar. **C)** Scatter plot comparing log_2_ fold-change values for rhCD40L-stimulated αDC1 (*x* axis) and PGE_2_-DC (*y* axis) relative to their respective unstimulated states. Genes are categorized as exclusively upregulated in αDC1 (pink), exclusively upregulated in PGE_2_-DC (orange), shared between both DC types (blue), divergently regulated (green), or not significant due to |log_2_FC| < 1 and/or adjusted *p* ≥ 0.01 (grey); representative genes are labeled. **D)** Volcano plot of DEGs (total = 14,476 genes) between rhCD40L-stimulated αDC1 and PGE_2_-DC; genes with |log_2_FC| > 1 and adjusted *p* < 0.01 are highlighted and representative genes are labeled. **E, F)** KEGG pathway enrichment analysis of DEGs upregulated in rhCD40L-treated αDC1 (**E**) or PGE_2_-DC (**F**) as compared to the other stimulated DC type. Bubble size represents the number of contributing genes, fill color represents -log_10_(adjusted *p*), and *x* axis indicates fold enrichment (note different scales between each panel). All DEGs were determined by MAST with regression of percent mitochondrial transcripts and participant; pathway analysis was conducted on all statistically significant DEGs with |log_2_FC| > 1 using pathfindR.

### αDC1 comprise phenotypically and transcriptionally distinct subpopulations at rest and after stimulation

In addition to the mature DC types differentially responding to rhCD40L, we observed that αDC1 alone consisted of three transcriptionally distinct subclusters in both the resting and rhCD40L-treated states (Figure 3A; Supplemental Figure S2). Each cluster of rhCD40L-stimulated αDC1 expressed a unique signature of cytokines and chemokines with divergent implications for T cell priming. The αDC1 in the designated cluster D highly express the genes encoding CCL17 (2.56 log_2_ fold change) and CCL19 (0.92 log_2_ fold change) compared to the other two clusters (Figure 3B, C). CCL17 is known to preferentially attract CCR4-expressing Th2 and T_reg_ cells, particularly in the context of tumor microenvironments (32,33), and CCL19 promotes the recruitment of multiple types of CCR7^+^ T cells and naïve T cells (34). Furthermore, cluster D also highly transcribed *IL2RA* (Figure 3B), which codes for the alpha subunit of the IL-2 receptor (CD25) and can be secreted to modulate IL-2 signaling (35). Cluster F exhibited increased expression of genes encoding the chemokines CCL3, CCL4, and CCL20, as well as TNF-α (*TNF*), and IL-23p19 (*IL23A*) (Figure 3B, D). These chemokines are known attractants of naïve CD8^+^ T and existing Th17 cells, while IL-23p19 combines with IL-12p40 to yield the functional form of the cytokine IL-23, a critical factor involved in the priming and differentiation of naïve CD4^+^ T cells into effector Th17 cells (36–39). Interestingly, we found that the 0.77% of αDC1 stimulated with rhCD40L transcribing both subunits of IL-12p70 (*IL12A* and *IL12B*) were primarily located in Cluster E (Figure 3E, G). Furthermore, the expression of *IL23A* and *IL12B* was restricted to a small percentage (1.42%) of cells located in cluster F (Figure 3F, H). Consistent with this transcriptional data, an IL-23 ELISpot revealed that only 0.80% of αDC1 stimulated with rhCD40L and 2.27% of αDC1 stimulated with J558 cells expressing CD40L secreted IL-23. Based on these transcriptional signatures, we propose the following predicted functional designations for these αDC1 subclusters: cluster E as the ‘IL-12p70-producing effector’ subset, cluster D as the ‘regulatory/Th2-attracting’ subset, and cluster F as the ‘inflammatory/Th17’ subset.

**Figure 3.**
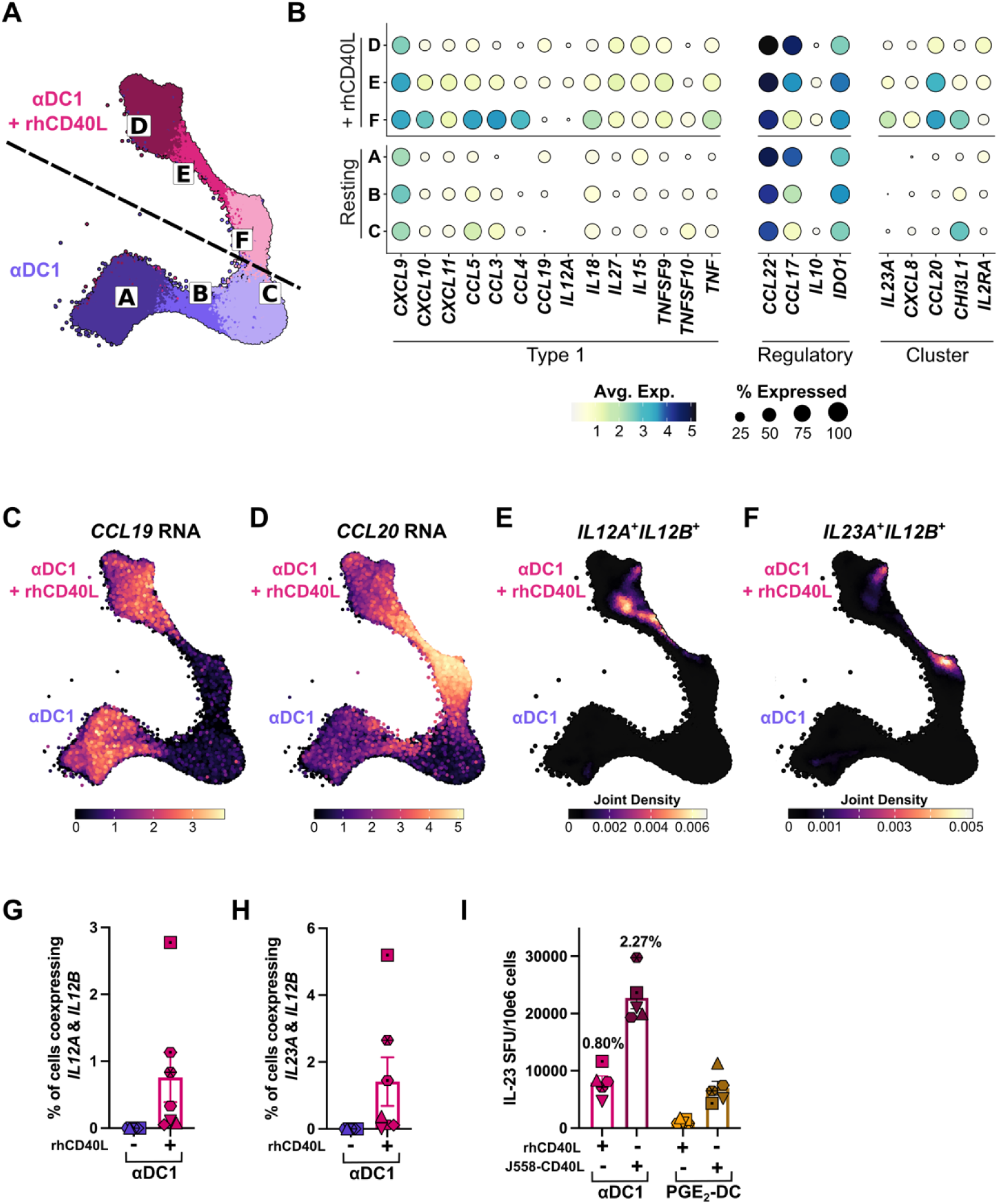
αDC1 resolve into three subpopulations with functionally distinct cytokine and chemokine transcription signatures. **A)** UMAP visualization of resting (subclusters A-C) and rh-CD40L-treated (subclusters D-F) αDC1 extracted from the overall UMAP in figure 2A (*n* = 7 participants). **B)** Dot plot representing relative expression of transcripts encoding selected cytokines and chemokines across the 6 αDC1 subclusters. These cytokines and chemokines are grouped into those promoting a type 1 response (Type 1), those supporting regulatory or Th2 responses (Regulatory), and others associated with individual subclusters (Cluster). Dot size represents the percentage of cells expressing the gene, and fill color indicates average scaled expression with darker shades indicating higher expression. **C, D)** Feature plots depicting *CCL19* (**C**) and *CCL20* (**D**) expression overlaid on the UMAP from panel A; lighter colors indicate higher scaled expression. **E, F)** Nebulosa kernel density estimation plot depicting the joint probability density of cells co-expressing the subunits of active IL-12p70, *IL12A* and *IL12B* (**E**), or active IL-23, *IL23A* and *IL12B* (**F**), across αDC1 subclusters in resting and rhCD40L-treated conditions. **G, H)** Percentage of αDC1 coexpressing *IL12A* (scaled expression > 1) and *IL12B* (scaled expression > 2.5) (**G**) or *IL23A* (scaled expression > 1.8) and *IL12B* (scaled expression > 2.5) (**H**) (*n*=7 participants). For both panels G and H, bivariate feature plots were used to determine an expression threshold identifying a discrete population with concurrent expression above background. **I**) The frequencies of IL-23-secreting αDC1 and PGE_2_-DC were determined via ELISpot after 12 hours of rhCD40L or J558-CD40L stimulation and expressed as SFU per 10^6^ cells. Labeled percentages indicate the mean percentage of IL-23-producing cells, and bars represent mean ± SEM. Low *n* precluded statistical analysis. Individual participant values are shown with unique symbols. All transcriptomic data represents *n* = 7 participants.

Given the heterogeneity present in both resting and rhCD40L-stimulated αDC1, we first investigated whether the transcriptional programs of each rhCD40L-treated subcluster were already present at baseline. We compared expression of key transcription factors, including stimulatory STATs (STAT1/2/4), IRFs, and canonical NF-κB subunits, as well as interferon-stimulated genes (ISGs), across paired resting and stimulated subpopulations (Figure 4A, B). Resting subclusters A, B, and C showed transcriptional profiles broadly consistent with their stimulated counterparts D, E, and F, respectively, suggesting that these functional states are established prior to CD40L exposure. Cluster B/E highly expresses *IRF8*, which promotes type 1 DC development and IL-12 production (40,41), providing further support for its characterization as the “IL-12p70-producing effector” subset. Cluster C/F exhibited high expression of stimulatory STATs, consistent with robust type I IFN responsiveness and autocrine IL-12/IL-23 signaling through STAT4 (42–44). During their generation for therapeutic use, αDC1 are not yet exposed to CD40L as part of their maturation strategy prior to delivery (45). Therefore, we next examined surface protein expression on the resting αDC1 subpopulations to identify candidates that could enable their isolation for optimal therapeutic potential. We included a panel of 42 oligomer-conjugated antibodies to surface proteins commonly present on APCs, allowing for direct comparison of the proteins expressed on the transcriptionally defined αDC1 subclusters. Surface protein expression profiles were similar between putative resting and stimulated counterpart clusters (Figure 4C), providing further support for pre- to post-rhCD40L-stimulation correspondence. Within the resting αDC1 subclusters A-C, we identified distinct protein expression patterns distinguishing each population (Figure 4D). Notably, cluster C highly expressed the mannose receptor CD206, the high-affinity Fc γ receptor CD64, and the tetraspanin CD81 (Figure 4D). Cluster A expressed relatively more integrin α4 (CD49d), CD162 (P-selectin glycoprotein ligand 1), the costimulatory molecule CD80, and the GM-CSF receptor CD116 (Figure 4D). Taken together, these data highlight the feasibility of isolating these distinct αDC1 subsets based on surface protein expression to optimize their therapeutic efficacy for various conditions.

**Figure 4.**
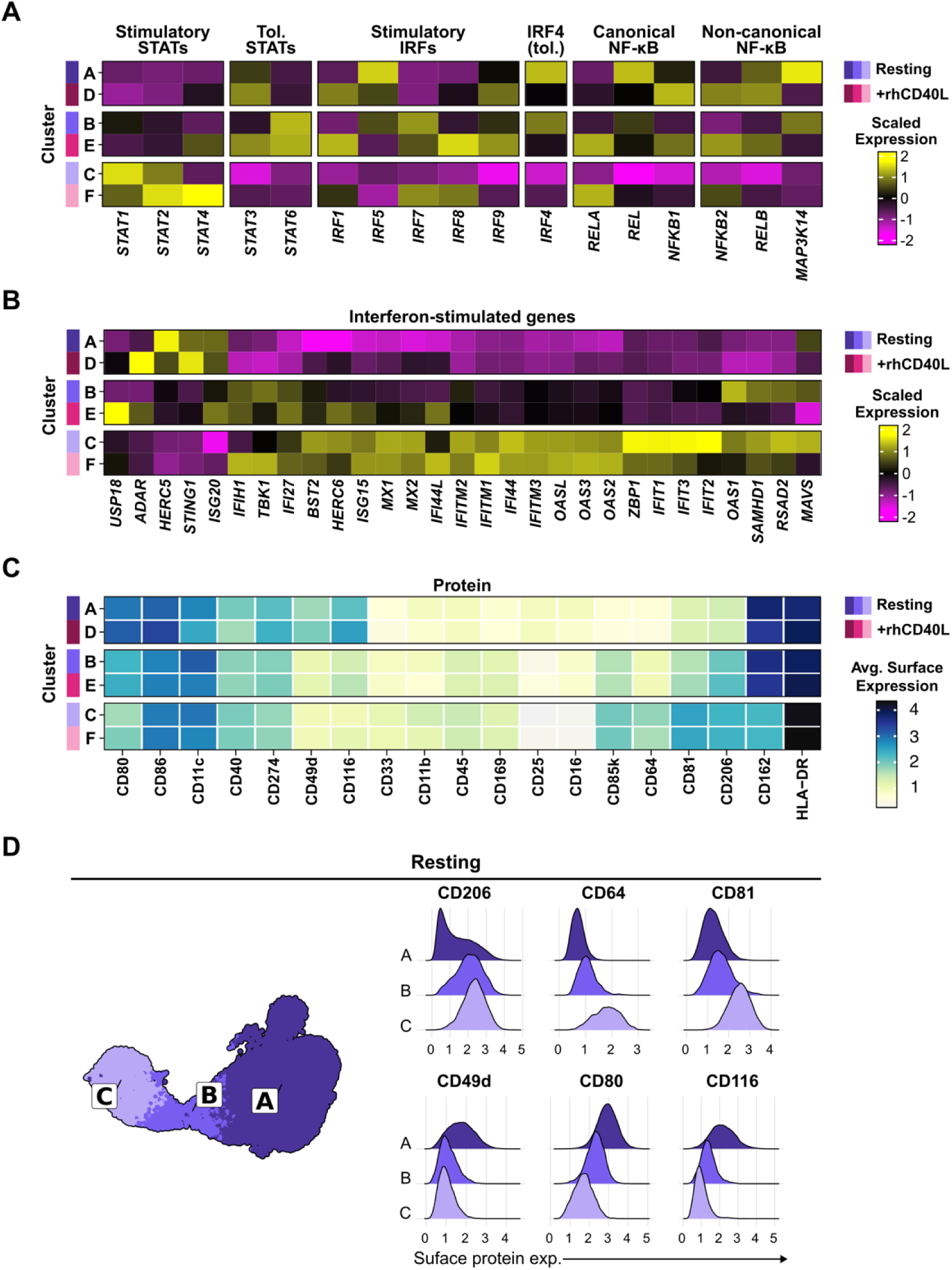
Resting and rhCD40L-stimulated αDC1 subclusters share transcriptional programs and are distinguishable by surface protein expression. **A, B)** Heatmap displaying average expression of transcripts encoding stimulatory and tolerogenic (tol.) transcription factors (panel **A**) and interferon-stimulated genes (panel **B**) across resting (A-C) and rh-CD40L-treated (D-F) αDC1 subclusters (*n* = 7 participants). Color scale represents scaled expression with yellow indicating higher relative expression and magenta indicating lower relative expression. Bar annotations denote cluster identity; purple bars represent resting subclusters and pink bars represent stimulated subclusters. **C)** Heatmap of average surface protein expression for 19 selected markers across both resting (A-C) and rh-CD40L-stimulated (D-F) αDC1 subclusters, derived from a 42-marker AbSeq panel. Color scale represents average expression with darker colors indicating higher expression. **D)** UMAP (left) of unstimulated αDC1 from *n* = 4 participants showing three subclusters (A-C) identified by weighted nearest-neighbor analysis of whole-transcriptome and surface protein data. Ridge plots (right) showing the distribution of CD206, CD49d, CD64, and CD80 surface protein expression across unstimulated αDC1 subclusters.

### Distribution among αDC1 subclusters varies between individuals

Because αDC1 prepared from different individuals produce widely varying amounts of important cytokines including IL-12p70, we investigated whether the relative abundance of the unique αDC1 subpopulations also exhibits interindividual heterogeneity. While both resting and rhCD40L-stimulated αDC1 generated from each of the study participants all consistently displayed the identified clusters A-F, the distribution among these clusters varied between the participants (Figure 5A, B). While 3 out of 4 individuals had low proportions of clusters C (6.4-17.7%) and F (1.4-5.4%), Participant 3 had a marked inflation of these populations with cluster C representing 39.8% of resting αDC1 and cluster F representing 25.6% of rh-CD40L-treated αDC1. Within individuals, the proportions of αDC1 in corresponding resting and stimulated subclusters remained stable across stimulation (Figure 5C). Importantly, this donor-intrinsic variation in subpopulation composition impacted bulk cytokine and chemokine production at the protein level. Principal component analysis of rhCD40L-induced cytokine and chemokine secretion separated participants in a manner that corresponded to their αDC1 cluster distributions, and the transcriptional signatures of each cluster drove this separation (Figure 5D). Participant 3, whose preparation was enriched for cluster C/F, demonstrated higher secretion of TNF-α, CCL3, and CCL4 (Figure 5D; Supplemental Figure S3), which all had upregulated transcript expression in cluster F in response to rhCD40L (Figure 3B). Taken together, these data suggest that resting subcluster proportions are a donor-intrinsic property that predicts the functional output of αDC1 preparations upon subsequent CD40L stimulation.

**Figure 5.**
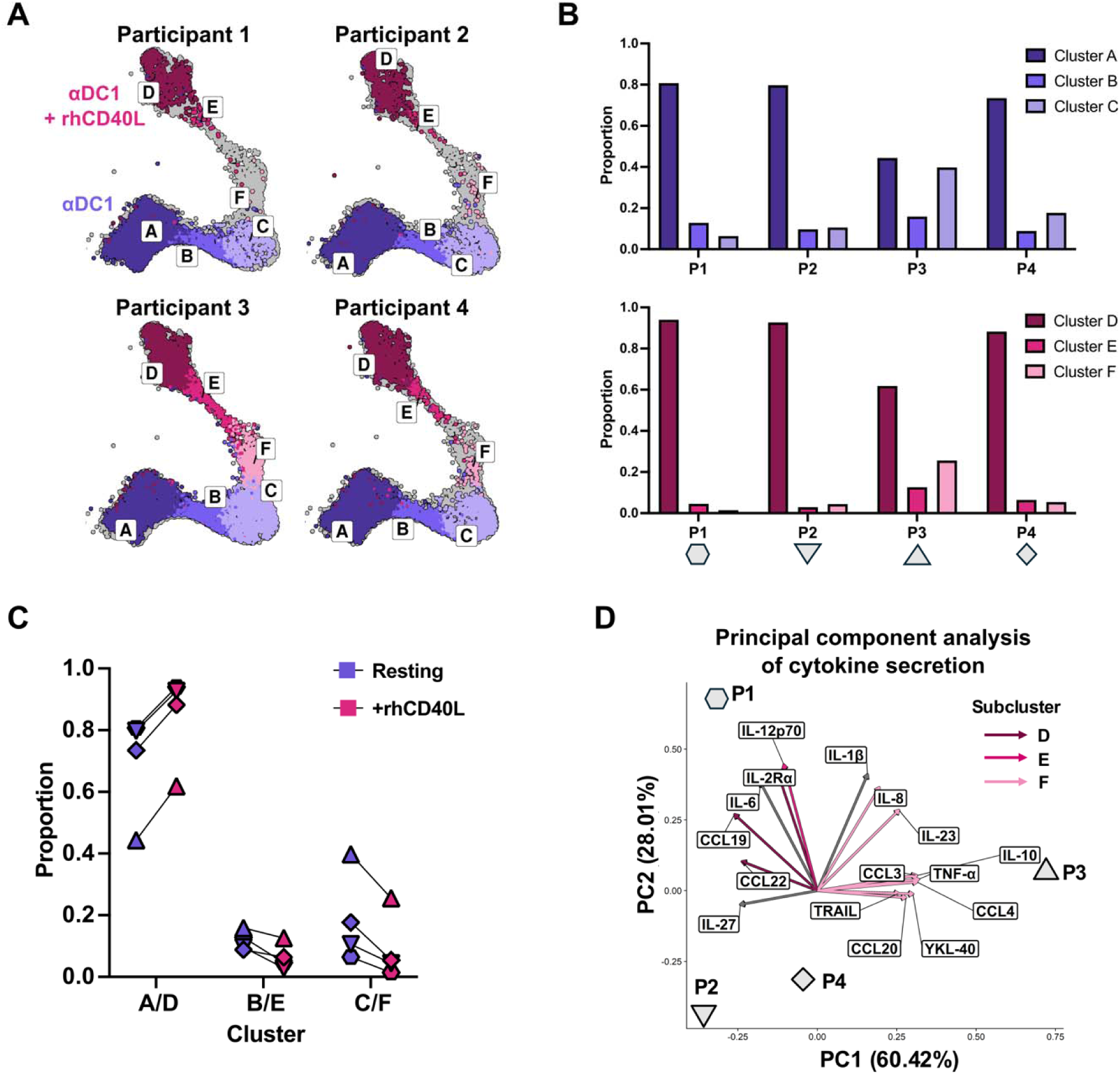
αDC1 subcluster proportions vary between individuals and predict bulk cytokine output. **A)** UMAP visualizations for four representative study participants exhibiting inter-individual variation in the distribution of αDC1 subclusters in both unstimulated and rhCD40L-treated conditions. **B)** Bar graphs showing the proportions of unstimulated subclusters A–C (upper) and rhCD40L-treated subclusters D–F (lower) for each participant (P1–P4). **C)** Paired comparisons of αDC1 subcluster proportions between unstimulated and rhCD40L-treated conditions for corresponding cluster pairs A/D, B/E, and C/F. Individual participant values are shown with unique shapes connected by lines to illustrate directional consistency across stimulation conditions (*n* = 4 participants). Formal significance testing was not performed, as the minimum achievable *p*-value for a paired test at this sample size precludes meaningful inference. **D)** Principal component analysis (PCA) biplot of bulk cytokine and chemokine concentrations measured by multiplex ELISA in supernatants from rhCD40L-stimulated αDC1 (total variance explained = 88.43%). Vector loadings are colored by associated αDC1 subcluster (D, E, or F), and individual participants are labeled (P1–P4).

## DISCUSSION

The αDC1 antigen presenting cell therapy platform, which has been widely explored in clinical trials for cancer as well as chronic HIV infection, was optimized for timely production of IL-12p70, a critical factor for driving cell-mediated immunity. However, our study clearly demonstrates that only a small fraction (0.5-2%) of αDC1 produce both subunits of IL-12p70 at the RNA and protein levels upon rhCD40L stimulation, despite releasing a much higher overall amount of this cytokine in comparison to PGE_2_-DC. Though this finding was unexpected given that αDC1 were developed for optimal and timely production of IL-12p70, it aligns well with emerging evidence of functional heterogeneity in well-defined DC subsets. Deak et al. identified murine ‘first responder’ conventional DC cycling through a transient active state that represent less than 5% of cells but generate over 70% of early inflammatory cytokine output (46), and Shalek et al. observed similar ‘precocious’ bone marrow-derived DC that quickly initiate an antiviral transcriptional program before later spreading it to other cells via paracrine IFN-β signaling (47). Similarly, a minority of plasmacytoid DC release type I interferon upon stimulation (48), and IL-12p70 production was limited to a subset of monocyte-derived DC matured by multiple stimulations (49). Our data extend this pattern to the clinically relevant monocyte-derived αDC1, identifying a discrete population (cluster E) characterized by expression of both *IL12A* and *IL12B* that is responsible for superior IL-12p70 production. While transcriptional noise and signaling thresholds may determine which cells respond at a particular time, stable epigenetic programming via DNA and histone methylation in the IL-12p70 subunit promoter regions also impacts a DC’s functional response (50,51). We therefore predict that cluster E cells are ‘effector αDC1’ representing a functionally superior subpopulation whose IL-12p70 production capacity may be a key determinant of the overall immunostimulatory potency of a given preparation, aligning our single-cell observations with the established clinical correlation between bulk IL-12p70 secretion and favorable outcomes in αDC1 vaccine trials (19,20,52,53).

The identification of other distinct αDC1 subpopulations with potentially antagonistic effects on T cell polarization has major therapeutic implications. For example, cluster D is characterized by high expression of *CCL17* and *CCL22*, which have been mechanistically linked to tumor immune evasion and correlated with relapse in breast cancer (54,55). This cytokine profile is strikingly similar to that of mature regulatory DC (mregDC) that recruit T_regs_ to establish a suppressive niche, dampening antigen trafficking to the draining lymph node and predicting worse outcomes in multiple cancers (56). Importantly, and consistent with previous findings, we show that PGE_2_-DC are much stronger producers of CCL22 compared to αDC1 at the protein level (Supplemental Figure S3), and CCL22 production by PGE_2_-DC has been shown to attract T_regs_ and therefore limit their utility in treating cancers and intracellular infections (57). The presence of this proposed ‘regulatory/Th2-attracting’ cluster D within αDC1 preparations suggests that this suppressive potential is not fully eliminated by the αDC1 maturation protocol, and depletion of cluster D cells represents a potential strategy for improving preparation quality. The cells within cluster F, which we propose designating as the ‘inflammatory/Th17’ subset due to their heightened expression of *IL23A*, present another therapeutic challenge. Though IL-23 production offers context-specific benefits, including Th17-mediated protection against fungal infections and potential utility in ovarian cancer (58,59) and HIV (60,61), this cytokine also stabilizes effector T_reg_s and can promote Th17-to-T_reg_ conversion within tumor microenvironments (62,63), indicating that cluster F enrichment may be counterproductive in most solid tumor applications. Taken together, the presence of individual populations within a single αDC1 preparation differentially biased toward effector (cluster E), suppressive (cluster D), and context-dependent inflammatory (cluster F) transcriptional programs suggests that therapeutic efficacy depends not only on total IL-12p70 secretion but on the balance among these subsets; therefore, independently targeting each subpopulation represents a rational strategy for optimizing αDC1 potency.

Our data demonstrate that the proportion of αDC1 subpopulations varies markedly between individuals, which provides a potential mechanism underlying inconsistent clinical outcomes observed in DC vaccine trials. A recent study analyzing the bulk transcriptomic profiles of autologous monocyte-derived DC vaccines used in a prostate cancer clinical trial identified three distinct maturation trajectories that varied among study participants, with a high type-I interferon response predicting positive clinical outcomes (64). Our identification of surface markers distinguishing αDC1 subsets at baseline offers a prospective strategy for enriching therapeutically favorable subclusters. Critically, because these subcluster proportions are detectable in the resting state and predict bulk cytokine output after CD40L stimulation, they could serve as a pre-administration biomarker to identify preparations likely to promote robust Th1 responses or to flag preparations enriched for potentially suppressive cluster D cells. The surface markers supporting this approach include CD49d, which is highly expressed on mature migration-competent DC, and Siglec-1/CD169, which is induced by strong type-I interferon signaling (65). Importantly, existing GMP-compliant DC isolation protocols underscore the technical feasibility of selecting for αDC1 subpopulations without using methods that alter function or phenotype (66,67).

We acknowledge several limitations of our study that warrant consideration. Because our single-cell analyses were performed at a fixed timepoint, we could not interrogate transitions between functional states similar to those observed in ‘first responder’ DC described in the report from Deak et al (46). This timepoint, however, does appear to capture the IL-12p70 response accurately given the concordance between the percentages of αDC1 transcribing both subunits at 4 hours and secreting protein following 18 hours of stimulation in the ELISpot assay. Although we cannot formally exclude the possibility that the identified subclusters represent transitional states, the presence of heterogeneity in resting αDC1, the stability of subcluster proportions within individuals across stimulation conditions, and the similar frequencies of IL-12p70-producing cells at both the 4 hour transcriptional and 18 hour protein analysis timepoints collectively suggest that these represent pre-committed functional states rather than transient intermediates. Additionally, our current study does not address whether the heterogeneity observed in the αDC1 preparations originates from monocyte subpopulations or variable sensitivity to αDC1-specific maturation factors at the immature DC stage. Previous work demonstrating that only classical monocytes differentiate into immature monocyte-derived DC when cultured with IL-4 and GM-CSF (68) and the relative homogeneity of the PGE_2_-DC suggest that the heterogeneity of αDC1 may instead arise from the immature DC response to polarizing stimuli. Multiomic analysis of other DC platforms designed for high IL-12p70 production, including those matured with CD40L and IFN-γ (20,69), could further elucidate how maturation cocktails shape heterogeneity in DC therapeutics. Furthermore, the ex vivo behavior of αDC1 subpopulations may not reflect their function in vivo once they are administered as therapeutics, where interactions with endogenous immune cells in tissue microenvironments could impact their performance. Finally, our findings are derived from a small number of healthy donors, and it remains to be determined whether the αDC1 heterogeneity and functional relationships described here extend to individuals living with HIV or cancer.

A retrospective analysis of cryopreserved samples from completed αDC1 vaccine trials could provide vital information about the relationship between subcluster distributions and clinical responses, potentially identifying the optimal balance for treating a particular disease. While we identified surface markers for αDC1 subsets, isolating these populations and comparing their ability to prime effector T cells and recruit distinct immune populations via established transwell migration assays (57) is essential to confirm that sorted αDC1 retain their predicted functional properties. This surface protein information can also be leveraged to establish a flow cytometry-based procedure for phenotyping αDC1 prepared for clinical use. Future studies should investigate whether enrichment of effector αDC1 (cluster E) improves Th1 priming efficacy and, ultimately, clinical outcomes. Additionally, the presence of a *CCL17*/*CCL22*-expressing cluster D subpopulation raises the possibility of depleting this subset as an alternative or complementary optimization strategy to effector αDC1 enrichment. Ultimately, the heterogeneity within αDC1 preparations identified in this study reveals multiple actionable avenues for further enhancing DC-based immunotherapies to maximize clinical benefit.

## METHODS

### Sex as a biological variable

Sex as a biological variable was not assessed in this study. Single-cell multiomics experiments were performed using a mixed cohort of leukocytes from donors without HIV collected by the Clinical Services Core D of the Rustbelt Center for AIDS Research (CFAR) (fully anonymized) and individuals without HIV who are participants of the Pittsburgh Clinical Research Site (CRS) of the MACS/WIHS Combined Cohort Study (MWCCS) (all assigned male at birth, reflecting site enrollment demographics). Given that donor sex was unavailable for a portion of the cohort, the remaining participants were all assigned male at birth, and the small sample size (*n* = 7), sex-stratified analyses were not feasible.

### Isolation of human primary monocytes

PBMC were isolated from donor leukapheresis products purchased from the Clinical Sciences Core D of the Rustbelt CFAR or whole blood obtained from men without HIV who are participants of the Pittsburgh CRS of the MWCCS via standard density gradient centrifugation using Lymphocyte Separation Medium (Corning Cat# 25–072-CV). Monocytes were then purified from PBMC using a positive selection human CD14 microbead kit (Miltenyi Biotec Cat# 130-097-052) according to the manufacturer’s protocol and cryopreserved until use.

### Generation of human monocyte-derived DC (MDC)

Immature MDC were generated by culturing peripheral blood isolated monocytes for 4 days at 37°C in IMDM (Gibco Cat# 12440-053) containing GM-CSF (1000 IU/mL; Sanofi Cat# NAC2004–5843-01) and IL-4 (1000 IU/mL; R&D Systems Cat# 204-1 L), supplemented with 10% heat-inactivated fetal bovine serum (GeminiBio Cat# 100-106-500) and 0.5% gentamicin (Gibco Cat# 15710-064) (cIMDM) plated in ultra-low attachment culture wells (Corning Cat# 3473). Previously described maturation cytokine cocktails containing either IFN-α (1000 IU/mL; Schering Corporation Cat# NDC:0085–1110-01), IFN-γ (1000 IU/mL; R&D Systems Cat# 285-1F), polyinosinic:polycytidylic acid (poly(I:C); 20 ng/mL; Sigma-Aldrich Cat# P9582-5MG), TNF-α (25 ng/mL; R&D Systems Cat# 210-TA), and IL-1β (10 ng/mL; R&D Systems #201-LB) or IL-6 (1000 U/mL; R&D Systems Cat# 206–1 L), prostaglandin E_2_ (PGE_2_; 2 μM; Sigma-Aldrich Cat# P6532-1MG), TNF-α (25 ng/mL), and IL-1β (10 ng/mL) were then added to the immature MDC to produce mature αDC1 or PGE_2_-DC, respectively (8,15). The cultures were incubated with the maturation factors for 48 hours before the cells were harvested and thoroughly washed for downstream use.

### Flow cytometry

Surface protein expression was analyzed by flow cytometry to phenotype MDC from n = 10 individuals. Cells were resuspended in FACS buffer consisting of 1X PBS (Cytiva Cat# SH30256.01), 0.5% BSA (Sigma Aldrich Cat# A7888-100G), and HEPES (10 mM; Gibco Cat#15-630-080) and blocked for 10 minutes at room temperature with human Fc block (50 µg/mL; BD Biosciences Cat# 564219, RRID:AB_2728082). The following antibodies were then added: CD83-PE (clone HB15A; Beckman Coulter Cat# IM2218U, RRID:AB_3741485), CD86-PE (clone HA5.2B7; Beckman Coulter Cat# IM2729U, RRID:AB_2917954), OX40L-PE (clone ik-1; BD Biosciences Cat# 558164, RRID:AB_647195), Siglec-1/CD169-PE (clone 7-239; BioLegend Cat# 346004, RRID:AB_2189029), and mouse IgG1-PE isotype control (clone MOPC-21; BD Biosciences Cat# 554680, RRID:AB_395506). After incubation for 20 minutes at room temperature, the cells were washed with FACS buffer and fixed with 2% PFA at 4°C overnight before analysis on a BD LSR Fortessa flow cytometer. Data were analyzed with FlowJo software (BD; multiple versions, RRID:SCR_008520) after quality control using PeacoQC (70).

### Functional characterization of mature DC

Mature DC were induced to secrete cytokines and chemokines as previously described (18). Briefly, harvested αDC1 and PGE_2_-DC were plated in a 96-well plate (2.5×10^4^ cells/well) and stimulated for 24 hours with either rhCD40L (1 µg/mL; Enzo Life Sciences Cat# ALX-522-110-C010), or CD40L-transfected J558 cells (J558-CD40L, RRID:CVCL_B7TP; 5.0×10^4^ cells/well) originally provided as a kind gift from Dr. Peter Lane (University of Birmingham, UK). Following stimulation, the supernatants were collected and tested for the presence of IL-12p70 by specific ELISA using the following kit reagents: rhIL-12 Standard (R&D Systems Cat# 219-IL-005), hIL-12 primary mAb (Thermo Fisher Scientific Cat# M122, RRID:AB_223556), biotin-labeled secondary hIL-12 mAb, (Thermo Fisher Scientific Cat# M121B, RRID:AB_223555), HRP-conjugated streptavidin (Thermo Scientific Cat# N100), TMB substrate solution (Thermo Scientific Cat# N301), and Stop Solution (Thermo Scientific Cat# N600). IL-23 concentration in supernatants was also measured by specific ELISA using a commercial kit according to the manufacturer’s instructions (MABTECH Cat# 3457-1HP, RRID:AB_3741486). Absorbance measurements for both ELISAs were obtained using a Thermo Scientific Varioskan LUX Multi-Mode Microplate Reader (RRID:SCR_026792). Supernatants were also analyzed for the presence of the following cytokines and chemokines using the multiplex Meso Scale Discovery (MSD) electrochemiluminescence platform: IL-12p70, IL-1β, IL-8, IL-23, IL-10, CCL3, TNF-α, CCL4, YKL-40, CCL20, TRAIL, IL-27, CCL22, CCL19, IL-6, and IL-2Rα. For MSD analysis, the supernatants were prepared according to manufacturer’s specifications, plated onto a custom U-plex plate, and analyzed on the MESO QuickPlex SQ 120 (RRID:SCR_020304). Protein concentrations derived from both the IL-12p70 ELISA and MSD analyses were normalized to ATP concentration measured with the CellTiter-Glo 2.0 assay (Promega Cat# G9241). 2.5×10^4^ cells were plated in a white 96-well optical-bottom plate (Thermo Fisher Scientific Cat# 165306) concurrently with plating for protein analysis. The CellTiter-Glo 2.0 assay was then immediately performed according to the manufacturer’s protocol with a standard curve and luminescence was read on a Thermo Scientific Varioskan LUX Multi-Mode Microplate Reader (RRID:SCR_026792).

### Quantification of cytokine-producing DC

The numbers of IL-12p70- and IL-23-producing mature DC were determined by ELISpot assay using IL-12/23 p40 capture antibodies (1 µg/well; clone MT86/221; MABTECH Cat# 3450-3-250, RRID:AB_907313) and IL-12p70-biotin (0.2 µg/well; clone MT704; MABTECH Cat# 3455-6-250) and IL-23p19-biotin (0.2 µg/well; clone MT155; MABTECH Cat# 3457-6-250, RRID:AB_10554340) detection antibodies, respectively. 1.25×10^4^ and 2.5×10^4^ harvested mature DC in AIM-V medium (Gibco Cat# 12055091) were added to each well of the coated ELISpot plate and incubated for 30 minutes at 37°C to allow the cells to adhere. The wells were then stimulated (as described above) with either rhCD40L, J558-CD40L or with AIM-V medium as an unstimulated control. After 18 hours, the plates were processed according to manufacturer’s specifications and analyzed on the AID iSpot EliSpot FluoroSpot Plate Reader (RRID:SCR_025287). Reported values are net responses compared to the resting control.

### Single cell multiomics analysis

αDC1 and PGE_2_-DC from *n* = 7 independent study participants were sampled for single cell multiomics analysis using the BD Rhapsody Single-Cell Analysis System (RRID:SCR_027096) based on their combined representation of the spectrum of CD40L-induced IL-12p70 production. The mature DC were harvested on culture day 6, washed to remove maturation cytokines, and replated in a fresh ultra-low attachment 24 well plate before stimulation with 1 µg/mL rhCD40L for 4 hours. The cells were then collected and labeled with condition-specific oligomer-conjugated multiplexing antibodies (BD Biosciences Cat# 633781, RRID:AB_2870299). For *n* = 3 participants, the multiplexed cells were stained with the following antibodies to analyze surface protein expression: CD19 (clone HIB19; BD Biosciences Cat# 940247, RRID:AB_2876128), CD274 (clone MIH1; BD Biosciences Cat# 940035, RRID:AB_2875926), CD3 (clone SK7; BD Biosciences Cat# 940000, RRID:AB_2875891), CD86 (clone 2331 (FUN-1); BD Biosciences Cat# 940025, RRID:AB_2875916), HLA-DR (clone G46-6; BD Biosciences Cat# 940010, RRID:AB_2875901), and CD169 (clone 7-239; BD Biosciences Cat# 940223, RRID:AB_2876104). For *n* = 4 participants, the multiplexed cells were then stained with the OMICS-One APC panel of oligomer-conjugated AbSeq antibodies (BD Biosciences Cat# 572435) and 12 additional AbSeq antibodies (see Table S1) to analyze surface protein expression. Barcoded cells were then mixed in equal parts before loading onto a BD Rhapsody high-throughput cartridge per manufacturer’s instructions for single cell capture using the BD Rhapsody express instrument. Quality control metrics were verified using the BD Rhapsody scanner. cDNA libraries of the whole transcriptome, multiplexing barcodes, and AbSeq barcodes were then prepared according to manufacturer’s specifications. Pooled libraries were submitted to the Emory Yerkes National Primate Research Center Genomics Core (RRID:SCR_026418) for sequencing on an Illumina NovaSeq 6000 (RRID:SCR_016387) or the UCLA Technology Center for Genomics and Bioinformatics (RRID:SCR_012204) for sequencing on an Illumina NovaSeq X (RRID:SCR_024569). Demultiplexing, initial cell calling, quality filtering, alignment, and annotation were performed using the BD Rhapsody Whole Transcriptome Analysis pipeline on the Seven Bridges Genomics platform (multiple versions, RRID:SCR_008308). The resulting Seurat objects were then imported into R Studio (R version 4.5.0, RRID:SCR_001905), and Seurat (version 5.0.3, RRID:SCR_016341) was used for further quality control (71). After removing multiplets and cells with mitochondrial reads >25%, *n* = 149,535 cells were included in the analysis. Samples were log normalized using Seurat::NormalizeData and the first 3 participants were batch-corrected using the RPCA method implemented in Seurat::IntegrateData. Because the average sequencing depth (83,368.42 reads/cell) was greater for these 3 participants compared to the others (48,083.84 reads/cell), the remaining 4 participants were reference-mapped using that integrated dataset as the reference before downstream analysis. After re-scaling the mapped data, differential gene expression analysis was performed using MAST (version 1.33.0) with regression of percent mitochondrial transcripts and participant as implemented in Seurat::FindMarkers. Cells were randomly downsampled for Supplemental Figure S1 for DEG analysis for computational efficiency. The data were plotted using the SCpubr (version 3.0.0; preprint citation), EnhancedVolcano (version 1.26.0, RRID:SCR_018931), ComplexHeatmap (version 2.24.1, RRID:SCR_017270), scCustomize (version 3.0.1, RRID:SCR_024675), and UpSetR (version 1.4.0, RRID:SCR_026112) packages (72–76). After filtering for DEGs with *p* adj < 0.01 and |log_2_FC| > 1, pathway enrichment analysis was conducted on those gene sets using the pathfindR package with the Biogrid PIN (version 2.6.0) (77). Proteomic data was normalized using centered log ratio as implemented in Seurat::NormalizeData.

### Statistics

Flow cytometry MFI data were assessed for normality by Q-Q plot inspection. Pairwise comparisons across DC maturation states were performed using Wilcoxon matched-pairs signed-ranks test with Bonferroni correction applied for multiple comparisons (significance threshold *p* < 0.0167; Figure 1B). IL-12p70 concentrations were log-transformed to achieve normality before comparison across DC type and stimulation condition using repeated measures two-way ANOVA with Šídák’s correction for multiple comparisons (Figure 1C). IL-12p70 ELISpot data were compared between DC types using Wilcoxon matched-pairs signed-ranks test (Figure 1D). All statistical analyses were performed in GraphPad Prism (version 11.0.0, RRID:SCR_002798). A *p* value < 0.05 was considered statistically significant unless otherwise noted.

### Study approval

Peripheral blood was collected from men without HIV who are participants at the Pittsburgh CRS of the MWCCS. Leukapheresis products collected from anonymized donors without HIV were purchased from the Clinical Sciences Core D of the Rustbelt CFAR. These studies were approved by the IRB at the University of Pittsburgh (STUDY21050072; STUDY22110116). All participants provided written informed consent. Experiments were conducted in accordance with local regulations and with approval from the IRB at the University of Pittsburgh (STUDY21080110).

## Data availability

Values for all data points in graphs are reported in the Supporting Data Values file. All deidentified raw and processed single cell RNA sequencing data generated in this study will be deposited in the GEO database (accession number pending). Code used for analysis will be available in a Zenodo repository upon publication.

## CONTRIBUTORS

AED contributed to experimental work, data collection and analysis, and writing of the manuscript; PEJS contributed to the experimental work, data analysis, and writing of the manuscript; HB contributed with technical assistance and data collection; JJM contributed to study design, methodology, and data analysis; BJCM contributed to study design and investigation; CRR contributed funding acquisition, resources, and investigation; PK contributed to data interpretation, writing, and editing; RBM contributed to study design, conceptualization, writing and editing, funding acquisition, resources, and supervision.

## ACKNOWLEDGEMENTS

The authors thank Alok V. Joglekar, Thomas E. Smithgall, Ronald Slomba, and Bowen Dong for helpful discussions. The authors would like to give special recognition to Chloé I. Charendoff and Dan MacDonald from Waters Biosciences (formerly BD Biosciences) for their extraordinary commitment to providing advice and technical assistance when needed. The authors would also like to acknowledge the UCLA Technology Center for Genomics & Bioinformatics for technical support with library sequencing. We also express our sincere gratitude and appreciation to the dedicated staff of the Pittsburgh CRS of the MWCCS, and most importantly to the Pittsburgh MWCCS CRS study participants for their generous donation of time and specimens.

