## Supplemental Figures S1-S3; Supplemental Table S1 for "Type 1-polarized DC immunotherapeutic contains heterogeneous populations with IL-12p70 production restricted to a rare subset"

**Table of contents:**

Supplemental Figure 1 (page 2; legend page 3)

Supplemental Figure 2 (page 4; legend page 5)

Supplemental Figure 3 (page 6)

Supplemental Table 1 (page 7)

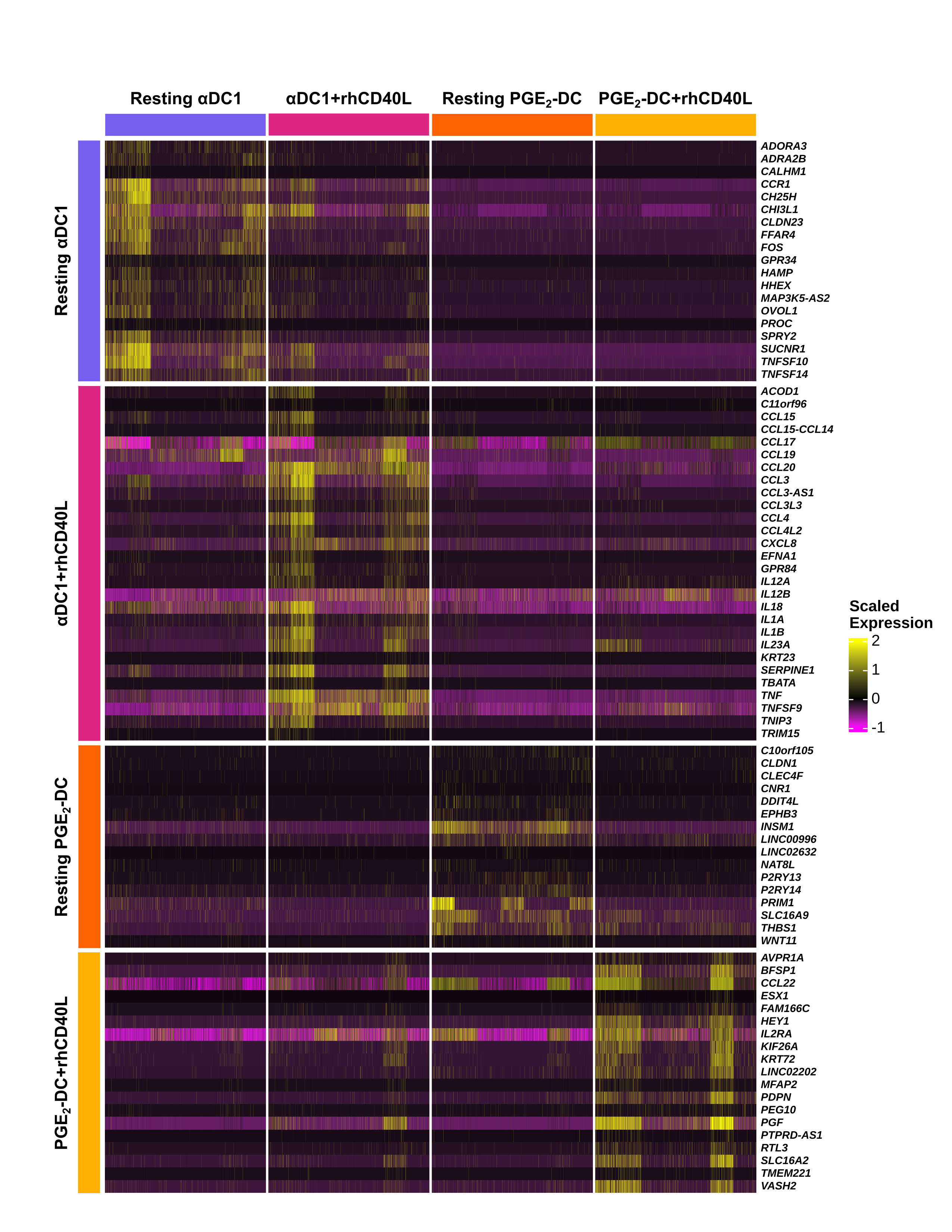

**Supplemental Figure S1: Differentially expressed genes across αDC1 and PGE_2_-DC ± rhCD40L stimulation.**

Heatmap displaying scaled expression of the top differentially expressed genes (DEGs) identified in each DC type both with and without rhCD40L stimulation (*n* = 7 participants). Equal numbers of cells were sampled from each participant within each condition for visualization. Columns represent individual cells grouped by condition, and rows represent individual genes. Genes are grouped by the condition in which they are most highly expressed, as indicated by the row annotations at left. DEGs were determined by running MAST with regression of percent mitochondrial transcripts and participant on a randomly downsampled set of 10,000 cells per condition.

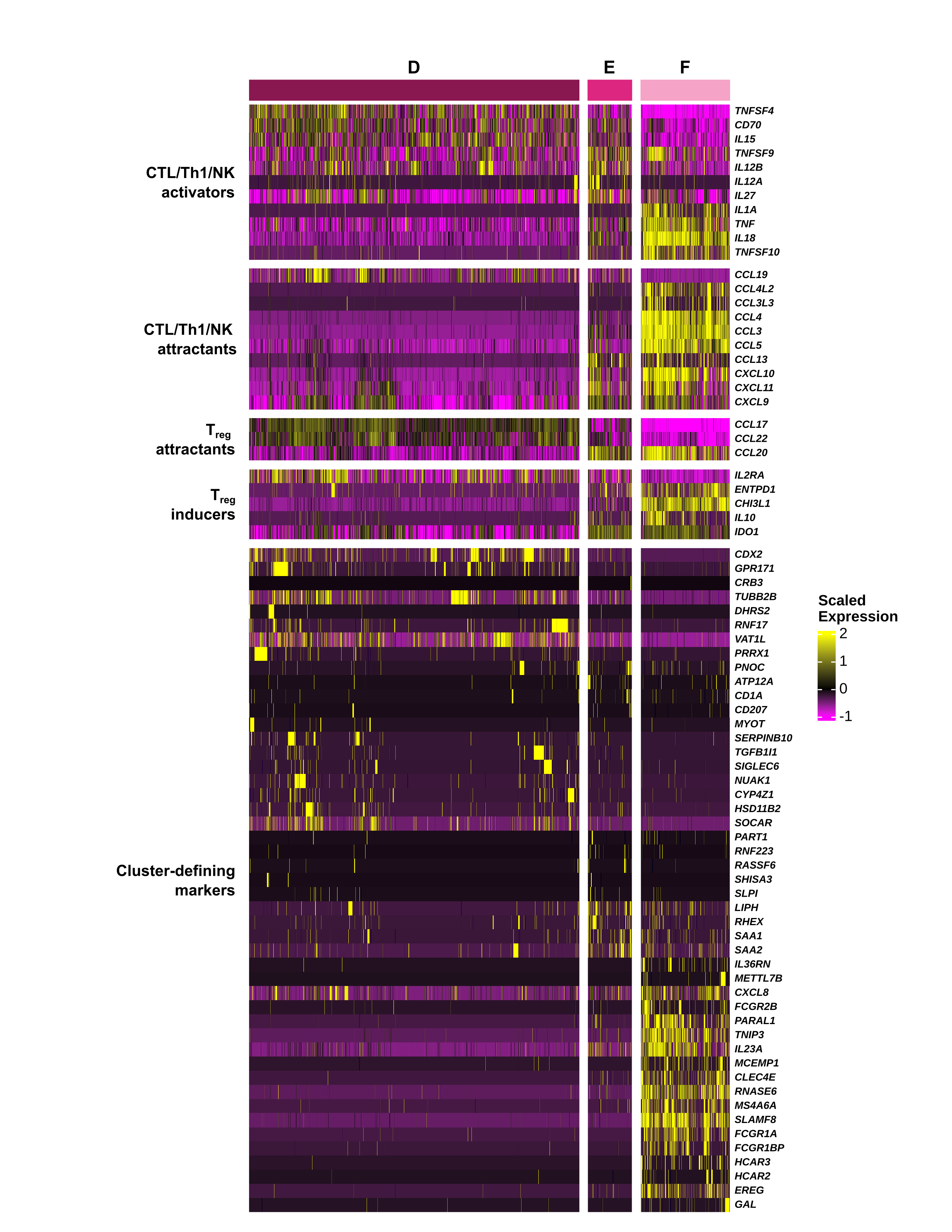

**Supplemental Figure S2: Differentially expressed genes across rhCD40L-stimulated αDC1 subclusters.**

Heatmap displaying scaled expression of differentially expressed genes (DEGs) identified in each rhCD40L-stimulated αDC1 subcluster (D, E, F) (*n* = 7 participants). Equal numbers of cells were sampled from each cluster for visualization. Columns represent individual cells grouped by subcluster; rows represent individual genes. Genes are grouped into CTL/Th1/NK cell activators or attractants, T_reg_ attractants or inducers, and remaining cluster-specific markers as indicated by the row annotations at left. DEGs were determined by running MAST with regression of percent mitochondrial transcripts and participant.

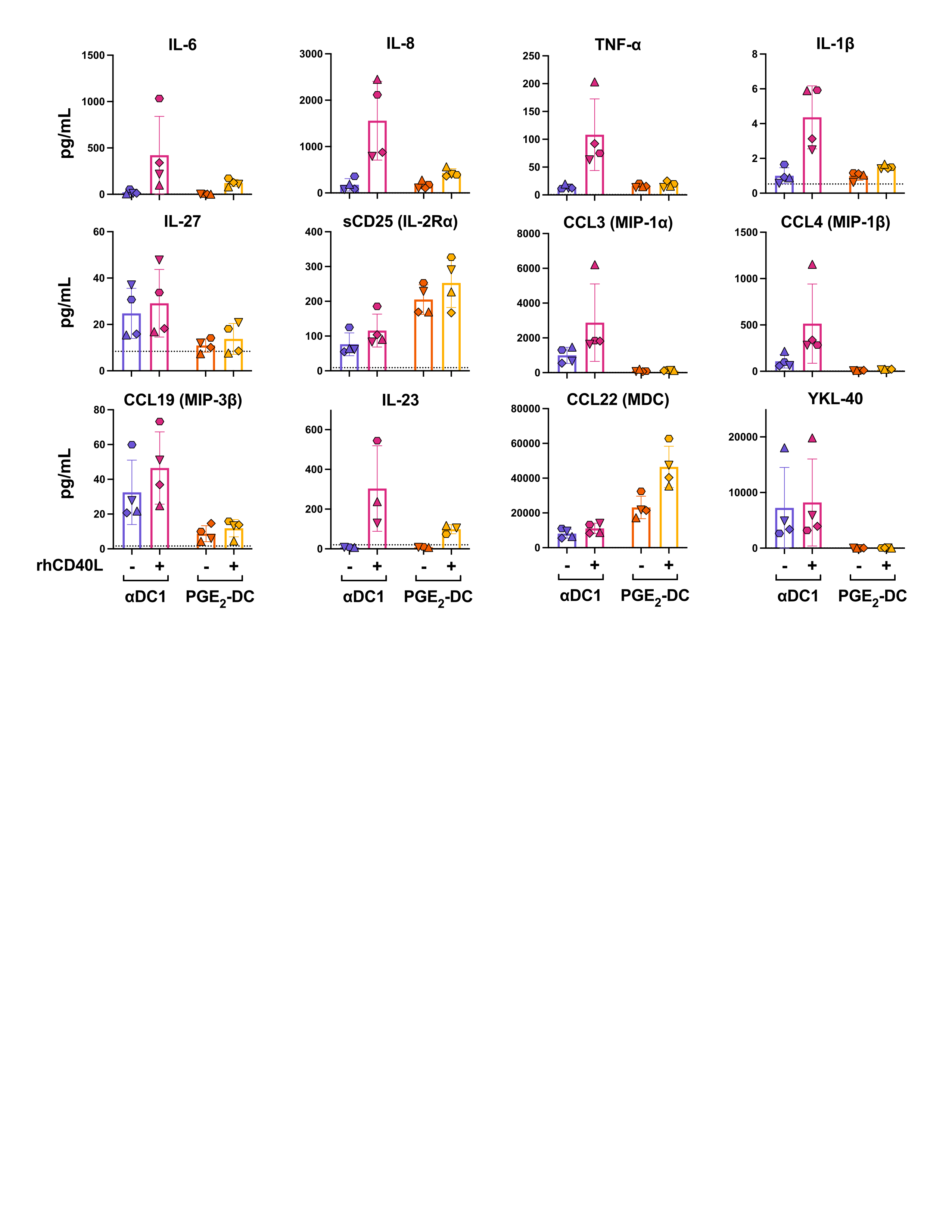
**Supplemental Figure S3. Multiplex analysis of cytokine and chemokine secretion by αDC1 and PGE₂-DC ± rhCD40L.**

Concentrations of IL-6, IL-8, TNF-α, IL-1β, IL-27, sCD25 (IL-2Rα), CCL3 (MIP-1α), CCL4 (MIP-1β), CCL19 (MIP-3β), CCL22 (MDC), and YKL-40 in supernatants collected from αDC1 and PGE_2_-DC following 24-hour stimulation with or without rhCD40L (1 µg/mL) were measured by multiplex electrochemiluminescence assay (Meso Scale Discovery) (*n* = 4 participants). IL-23 concentration was determined after the same stimulation by ELISA (*n* = 3 overlapping participants). Values were normalized to cellular ATP content. Dashed lines indicate assay lower limits of detection, and values below that limit are represented as half of the lower limit of detection for analysis purposes. Bars represent mean ± SD; individual participant values are shown with unique symbols. Low *n* precluded statistical analysis.

**Supplemental table 1. AbSeq antibodies used for CITE-seq proteomic analysis**

| **Target** | **Clone** | **BD cat #** | **RRID** |
| --- | --- | --- | --- |
| CD1c | F10/21A3 | OMICS-One APC/Myeloid-Cell Protein Panel (572435) | AB_2875974 |
| CD11c | B-LY6 |  | AB_2875915 |
| CD11b | M1/70 |  | AB_2875899 |
| CD14 | MPHIP9 |  | AB_2875896 |
| CD15 | W6D3 |  | AB_2876152 |
| CD16 | 3G8 |  | AB_2875897 |
| CD32 | FLI8.26 |  | AB_2875960 |
| CD33 | WM53 |  | AB_2875922 |
| CD36 | CLB-IVC7 |  | AB_2876105 |
| CD40 | 5C3 |  | AB_2875940 |
| CD45 | HI30 |  | AB_2875893 |
| CD64 | 10.1 |  | AB_2875914 |
| CD85k | ZM3.8 |  | AB_2876140 |
| CD80 | L307.4 |  | AB_2875927 |
| CD116 | hGMCSFR-M1 |  | AB_2876186 |
| CD103 | BER-ACT8 |  | AB_2875958 |
| CD115 (CSF1R) | 9-4D2-1E4 |  | AB_2876106 |
| CD123 | 7G3 |  | AB_2875911 |
| CD141 | 1A4 |  | AB_2875970 |
| CD162 | KPL-1 |  | AB_2876108 |
| CD163 | GHI/61 |  | AB_2875949 |
| CD169 | 7-239 |  | AB_2876104 |
| CD192 (CCR2) | LS132.1D9 |  | AB_2876163 |
| CD195 (CCR5) | 2D7/CCR5 |  | AB_2875941 |
| CD206 | 19.2 |  | AB_2875959 |
| CD273 | MIH18 |  | AB_2875962 |
| CD274 | MIH1 |  | AB_2875926 |
| HLA-DR | G46-6 |  | AB_2876102 |
| FCeR1a | AER-37 |  | AB_2875901 |
| VISTA | MIH65.rMAb |  | AB_2876339 |
| CD137 | 4B4-1 | 940055 | AB_2875946 |
| CD49d | 9F10 | 940059 | AB_2875950 |
| CD63 | H5C6 | 940243 | AB_2876124 |
| CD81 | JS-81 | 940052 | AB_2875943 |
| CD18 | 6.7 | 940086 | AB_2875977 |
| CD85j | GHI/75 | 940249 | AB_2876130 |
| HLA-DQ | TU169 | 940233 | AB_2876114 |
| CD89 | A59 | 940277 | AB_2876155 |
| CCR7 | L1-A | 940394 | AB_2876258 |
| CD25 | 2A3 | 940009 | AB_2875900 |
| CD83 | HB15e | 940054 | AB_2875945 |
| CD86 | 2331 (FUN-1) | 940025 | AB_2875916 |
